# Deep Learning-driven Automatic Nuclei Segmentation of Label-free Live Cell Chromatin-sensitive Partial Wave Spectroscopic Microscopy Imaging

**DOI:** 10.1101/2024.08.20.608885

**Authors:** MD Shahin Alom, Ali Daneshkhah, Nicolas Acosta, Nick Anthony, Emily Pujadas Liwag, Vadim Backman, Sunil Kumar Gaire

## Abstract

Chromatin-sensitive Partial Wave Spectroscopic (csPWS) microscopy offers a non-invasive glimpse into the mass density distribution of cellular structures at the nanoscale, leveraging the spectroscopic information. Such capability allows us to analyze the chromatin structure and organization and the global transcriptional state of the cell nuclei for the study of its role in carcinogenesis. Accurate segmentation of the nuclei in csPWS microscopy images is an essential step in isolating them for further analysis. However, manual segmentation is error-prone, biased, time-consuming, and laborious, resulting in disrupted nuclear boundaries with partial or over-segmentation. Here, we present an innovative deep-learning-driven approach to automate the accurate nuclei segmentation of label-free live cell csPWS microscopy imaging data. Our approach, csPWS-seg, harnesses the Convolutional Neural Networks-based U-Net model with an attention mechanism to automate the accurate cell nuclei segmentation of csPWS microscopy images. We leveraged the structural, physical, and biological differences between the cytoplasm, nucleus, and nuclear periphery to construct three distinct csPWS feature images for nucleus segmentation. Using these images of HCT116 cells, csPWS-seg achieved superior performance with a median Intersection over Union (IoU) of 0.80 and a Dice Similarity Coefficient (DSC) score of 0.88. The csPWS-seg overcame the segmentation performance over the baseline U-Net model and another attention-based model, SE-U-Net, marking a significant improvement in segmentation accuracy. Further, we analyzed the performance of our proposed model with four loss functions: binary cross-entropy loss, focal loss, dice loss, and Jaccard loss. The csPWS-seg with focal loss provided the best results compared to other loss functions. The automatic and accurate nuclei segmentation offered by the csPWS-seg not only automates, accelerates, and streamlines csPWS data analysis but also enhances the reliability of subsequent chromatin analysis research, paving the way for more accurate diagnostics, treatment, and understanding of cellular mechanisms for carcinogenesis.

## 1. Introduction

Advancements in medical imaging and computational analysis have been pivotal in enhancing our understanding of cellular mechanisms, particularly in the field of oncology [1]. The development of Chromatin-sensitive Partial Wave Spectroscopic Microscopy (csPWS) [2, 3] marked a significant milestone, enabling the nanosensing of cellular or intracellular structures with unprecedented details. This has opened new avenues for cancer research, allowing the early detection of malignancies through the analysis of chromatin dynamics, compaction, and topology within cell nuclei [4]. The cell nuclear architecture is mainly regulated by the interplay between the dynamic 3D chromatin organization and the distribution of genetic material. This chromatin structure is recognized as a common denominator for the detection of cancer-promoting status in the nucleus [5].

Chromatin regulates gene expression across multiple scales. These include the nucleosomal scale (∼10 nm, <1 kb genomic scale), involving histone and DNA modifications like acetylation and methylation, influencing which genes are active or silent; genome topology (>100 nm, >100 kb), involving chromatin loops, topologically associated domains (TADs), and A/B compartments, determining which parts of the genome are accessible to the cellular machinery that reads genetic information; and chromatin conformation within packing domains (PDs), spanning kilobase to megabase scales, influencing which genes are expressed over multiple cell generations [6,7]. Further, chromatin in the nucleus is known to be organized into several thousand PDs - compact regions that maintain heritability across cell divisions - varying in size (averaging 250 kb). These domains exhibit a chromatin volume concentration (CVC) of approximately 0.35 and demonstrate mass-fractal intradomain chromatin conformation with the genomic length *N*, scaling with the radius of the occupied physical volume *R* as *N* ∝ *R*^*D*^ where experimentally measured *D* values range from 5/3 to 3 across packing domains [3,8]. Thus, packing scaling *D* provides insight into how nanoscale materials are organized within PDs in a fixed volume.

The small size of the chromatin chain (∼20 nm) within chromatin PDs prevents the assessment of intradomain chromatin conformation using diffraction-limited optical microscopy (∼200 nm resolution). Despite extensive technological advancement in this area, the current understanding of chromatin organization is still limited for this nanoscale resolution. For imaging at this length scale, techniques such as chromatin conformation capture methods (chromosome conformation capture (3C), high-throughput 3C (Hi-C), circular chromosome conformation capture (4C), etc.) or electron microscopy are extensively used, which involve intensive sample preparation and fixation method and prevent temporal analysis of live-cell [4]. Recently, super-resolution microscopy techniques emerged as methods to study the spatial organization of higher-order chromatin structures beyond the diffraction limit resolution [7,9]. However, these techniques are limited to fixed cells and require exogenous labeling of cellular samples, which introduces additional width to the surface of cellular structures due to the physical dimension of dyes, introducing potential artifacts. These limitations become particularly significant when imaging the smaller size of the chromatin chain or DNA. These limitations may be addressed by label-free nanoimaging techniques, which can offer unbiased and accurate insights into cellular nanostructures and their dynamics. By relying on the intrinsic properties of cells, label-free techniques preserve the natural state of live cells, enabling the monitoring processes in real-time without the potential distortions introduced by dyes or stains. The csPWS microscopy is one such label-free optical non-invasive nanosensing technique that characterizes the relationship between genomic and physical scales of PDs inside live-cell nuclei using endogenous contrast without any fluorescent staining [2]. It measures variations in the interference spectrum of backscattered light from label-free samples and is sensitive to the PDs scaling, *D*, of chromatin chain conformation within PDs and its volume fraction.

Computational analysis of chromatin packing, dynamics, and topology within the live-cell nuclei in csPWS images requires isolating nuclei from the cytoplasmic and background structures. Thus, robust, reliable, and fast segmentation of all nuclei in csPWS images is the critical step for such assessment. This is especially true for oncology, where the segmented nuclei analysis enables researchers to discover chromatin structure modification within the cancer cell, allowing them to study the defense mechanisms against the treatment, which will be a very effective approach. Recently, csPWS has been utilized in human studies for the early detection of deadly cancers, including lung and colorectal cancer, and has demonstrated high efficacy in detecting Stage-I lung cancer (AUC = 0.92 in Site 1 and 0.81 in Site 2) [3] and advanced adenoma (AUC = 0.90) [10]. Furthermore, cancer heterogeneity poses a significant challenge to cancer research, while automated segmentation facilitates the gathering and evaluation of a large number of nuclear features from extensive cell populations. This approach generates ample data for statistical modeling, facilitating the identification of features pertinent to cancer cells.

The existing manual nuclei segmentation using the traditional methods is markedly labor-intensive and time-consuming. An expert user can draw all nucleus regions of interest (ROIs) in a csPWS image of size 1024 × 1024 pixels with 15 to 20 cells in approximately 10 to 15 minutes (∼ 45 sec/nucleus). Additionally, traditional manual segmentation is susceptible to boundary error, individual user selection bias, and misrepresentation, resulting in disrupted nuclear boundaries with partial or over-segmentation. Such limitations not only impede the throughput of nuclei chromatin analysis but also affect the potential accuracy of cancer diagnostics and overall progress within the field. Additionally, spatial segmentation of the nucleus has biological importance, and an accurate periphery analysis is crucial in cancer research. For example, in eukaryotic cells, the heterochromatin-enriched structures tend to localize toward the periphery, which highlights the importance of accurate detection of nucleus boundaries for a periphery analysis [11]. Besides that, the complexity of label-free imaging, such as lower resolution, lower signal-to-noise ratio, overlapping spectral features, heterogeneity in cell shapes and sizes, non-uniform intensity distributions of the same cell tissue, boundary ambiguity, etc., adds an extra challenge for accurate nuclei segmentation compared to its labeled counterparts [12].

Recently, deep learning (DL) emerged as an effective technique to analyze visual imagery tasks by automatically extracting features through convolution and filters from images instead of manual feature extraction. This automatic extraction process makes DL models highly accurate for several computer vision tasks. Image segmentation, where each pixel of an image is classified and grouped together based on similar attributes and features, such as color, intensity, or texture, is one of such fields that greatly benefited from DL and has been successfully showing state-of-the-art performance in various application areas such as medical imaging [13-15], autonomous vehicles [16,17], and so on.

Here, we introduce csPWS-seg, a computational strategy using DL for the automatic, accelerated, and accurate nuclei segmentation of the label-free csPWS images. csPWS-seg uses a gated channel transformation-based attention mechanism [18] integrated with baseline U-Net [19] (GCT-U-Net [20]) that provides accurate feature extraction, leading to automation of the accurate nuclei segmentation for csPWS microscopy imaging data. The attention mechanism in GCT-U-Net extracts more relevant features, contributing to a more accurate and improved nuclei segmentation by removing redundant and unnecessary features from the input images. Further, we analyzed the performance of our model on different kinds of segmentation loss functions. This analysis is performed on distribution-based segmentation loss functions, such as binary cross entropy loss [21] and focal loss [22,23], and the region-based loss function, such as dice loss [24] and Jaccard loss [25]. Finally, we compare our model’s performance with the baseline U-Net model and another attention-based model, SE-U-Net [26]. The csPWS-seg with focal loss provides the best performance among all models in live cell HCT116-cell line datasets captured using csPWS microscopy.

## 2. Material and Methods

### 2.1 csPWS microscopy and feature images

Conventional visible spectrum microscopy is limited to imaging of structures larger than 200 nm. csPWS, as a nanosensing method, detects chromatin packing domain changes in cells and, in doing so, distinguishes histologically normal cells potentially harboring cancer signatures. It sandwiches the nucleus between glass and media, capturing high-magnification, spectrally resolved images (450 - 700 nm) to analyze chromatin structure at sub-diffractional scales. The mechanism of csPWS microscopy is described in [3, 4, 9]. In brief, for a given location ***r*** within the cell, the refractive index (RI) *n* is proportional to the macromolecular density (*ρ*) of proteins, DNA, and RNA. This can be mathematically represented as *n*(***r***) = *n*_*water*_ + *αρ*(***r***) where *α* is the RI increment, which remains nearly constant for most macromolecules [3, 28]. For csPWS, a cell is nearly matched to the RI of media on the surface and mismatched to the other (glass).

For every pixel, csPWS measures the interference spectrum resulting from the interaction of the reference wave with scattered light across all RI fluctuations within a defined coherence volume. This volume is determined by spatial coherence in the lateral plane and longitudinal depth of field. Based on Parseval’s theorem, the standard deviation of the interference spectra (Σ) is proportional to the Fourier transform of the autocorrelation function (ACF) of *ρ*(***r***) integrated over the Fourier transform of the coherence volume. Σ is a measure of spatial variations of macromolecular density with subdiffractional sensitivity, sensing the length scales of 20-200 nm [29]. Given coherence volume centered around an intranuclear location, spectral bandwidth, N.A. values, nuclear thickness, and average domains’ CVC and genomic size, a genomic computational method [27] is utilized to translate Σ into csPWS packing scaling *D*, highlighting chromatin PDs upregulation.

The Σ image provides rich information on changes in mass distribution within the cell, which can be utilized for segmenting the nucleus from the cytoplasm. To enhance the accuracy and robustness of our AI-enhanced segmentation approach, apart from Σ image, we leveraged additional data provided by csPWS microscopy, including a brightfield (BF) image, single wavelength (SW) image, and reflectance image and evaluated their performance [30]. The BF image was obtained via brightfield microscopy, where white light illuminates the sample, and the image is formed by the light transmitted through the sample. The reflectance image was created by calculating the mean reflectance signal for each pixel across the entire spectrum. The SW image indicates the sample reflectance at 550 nm and was calculated by dividing the raw image of the sample at 550 nm by the raw image of a blank reference sample at 550 nm after both images had been compensated to account for internal reflections in the microscope objective. For our goal of isolating nuclei ranging from 1 micron to 5 microns (Buccal cells [31], Cancer cell lines [32]) from cytoplasm sized between a few to several hundred nanometers, using a wavelength of 550 nm appears to be an ideal choice. Biological samples exhibit a moderate scattering coefficient at 550 nm, while light absorption is relatively low [33], particularly in samples with high water content, such as cancer cell lines. At 550 nm, interference is moderately pronounced, leading to distinct constructive or destructive interference patterns based on height variations when light reflects off different layers on the nucleus surface (Interface Effect) in the SW image. Variations in the optical path length (OPL) from the reference surface to the scattering surfaces in the nucleus and cytoplasm induce phase differences in the reflected light, which results in a clearly visible interference pattern. This interference pattern has a distinct appearance in the nucleus, where shorter sub-micron OPLs are dominant, and the cytoplasm, where longer OPLs are dominant. This enables us to obtain characterizing information that helps to distinguish the nucleus from the cytoplasm.

### 2.2 csPWS Instrumentation

The live-cell csPWS instrumentation system has been previously described in reference [2,3]. In brief, it is built into a commercial inverted microscope (Leica, DMIRB, Wetzlar, Germany) equipped with a Hamamatsu Image Electron Multiplying Charge-Coupled Device camera (C9100-13, Hamamatsu City, Japan) coupled to a liquid crystal tunable filter (CRi, Woburn, MA, USA) to collect spectrally resolved Monochromatic images of the backscattered light between 500 and 700 nm with 1 nm step size. Further, broadband illumination is provided by an Xcite-120 LED lamp (Excelitas, Pittsburgh, PA, USA) supported by a tunable spectral collection filter. The sample is illuminated with a low N.A. of 0.52, whereas the collected backscattered light is limited by a high N.A. oil-immersion objective (100×, N.A. = 1.4). This yields a 3D data cube, I(*λ*, x, y), where *λ* is the wavelength and (x, y) correspond to pixel positions across a 10,000-λ^2^ field of view, allowing multiple cells (1–100, depending on the eukaryotic cell line) to be imaged simultaneously. Acquisition of the full interference spectra for spectral analysis takes under 17s, with each wavelength collection produced from < 100 ms exposures.

### 2.3 Cell Culture

HCT116 cells (ATCC #CCL-247) were cultured in McCoy’s 5A Modified Medium (#16600– 082, Thermo Fisher Scientific, Waltham, MA, USA) supplemented with 10% FBS (#16000– 044, Thermo Fisher Scientific, Waltham, MA, USA) and penicillin-streptomycin (100 μg/ml; #15140–122, Thermo Fisher Scientific, Waltham, MA, USA). All cells were maintained at 37 °C with 5% CO2 under the recommended conditions and were kept between passages 5 and 20. Cells were allowed at least 24 hours to re-adhere and recover from trypsin-induced detachment. Imaging was performed when the surface confluence of the dish was between 20% and 70%. Prior to starting perturbation experiments, all cells were tested for mycoplasma contamination (ATCC #30-1012K) and tested negative results. In this study, HCT116 cells were subjected to either mock treatment or drug treatment. For the mock-treated cells, the media was replaced after 24 hours. For the drug-treated cells, the media was replaced with a solution containing 15 μmol/L oxaliplatin, and the cells were treated for 48 hours.

### 2.4 csPWS Image Collection

The csPWS microscope was controlled via custom software with a graphical user interface (GUI). The imaging procedure began by scanning the whole glass bottom dish (well size 14 mm, #1.5 glass) using a 100× liquid objective in media for live cells and phosphate buffer saline (PBS) for fixed cells, selecting the cells for imaging by a trained user, and adjusting the focus for each image. The fluorescent imaging and/or csPWS spectral acquisition was performed with the cells in a liquid medium using a liquid-dipping 100× optical objective (Nikon, Melville, NY, USA). The csPWS acquisition software automatically acquired the spectral data for selected cells, and the analysis software rapidly generated the processed spectral data.

### 2.5 Deep Learning

#### 2.5.1 Training data

The training datasets for the DL are the set of csPWS microscopy images (input) and its nuclei-segmented images (labeled ground-truth). We generated 4 different training datasets using (i) BF image only, (ii) Σ image only, (iii) SW image only, and (iv) collections of BF, Σ and SW images. The labeled ground-truth datasets were obtained by manually segmenting each and every nucleus from all training data using the PWSpy GUI software module [30] by a postdoctoral-level expert with several years of experience in csPWS microscopy. Nuclei of all shapes and sizes were included. We tried to minimize the variation from manual selection bias, such as smoothing of the border lines due to shaking of the hand and modeling by ellipse-like equations with a small degree of freedom. Carefully taking the references from all three csPWS feature input images, an accurate ROI segmentation dataset for nuclei was prepared. Such careful and high-quality labeling is necessary to improve the model performance by reducing the error due to confounding factors of noisy shape, enabling all kinds of characteristic features to be learned, and reducing the false ROI detection. We prepared and trained three separate models with 210 csPWS images (BF, SW, and Σ models separately, each having approximately 5000 nuclei samples) of size 1024 × 1024 pixels from each of the 3 feature images. For the last model, a total of 630 csPWS images (collection of BF, SW, and Σ images with ∼ 15000 nuclei) were used. A sample training feature images with manually segmented ground truth are shown in Fig. 1. Further, to avoid overfitting, enough training datasets were generated by (i) performing data augmentation, such as horizontal & vertical flipping and 10% zooming each image, and (ii) dividing each image into non-overlapping patches of small size (e.g., we used 512 × 512 pixel size).

**Fig. 1.**
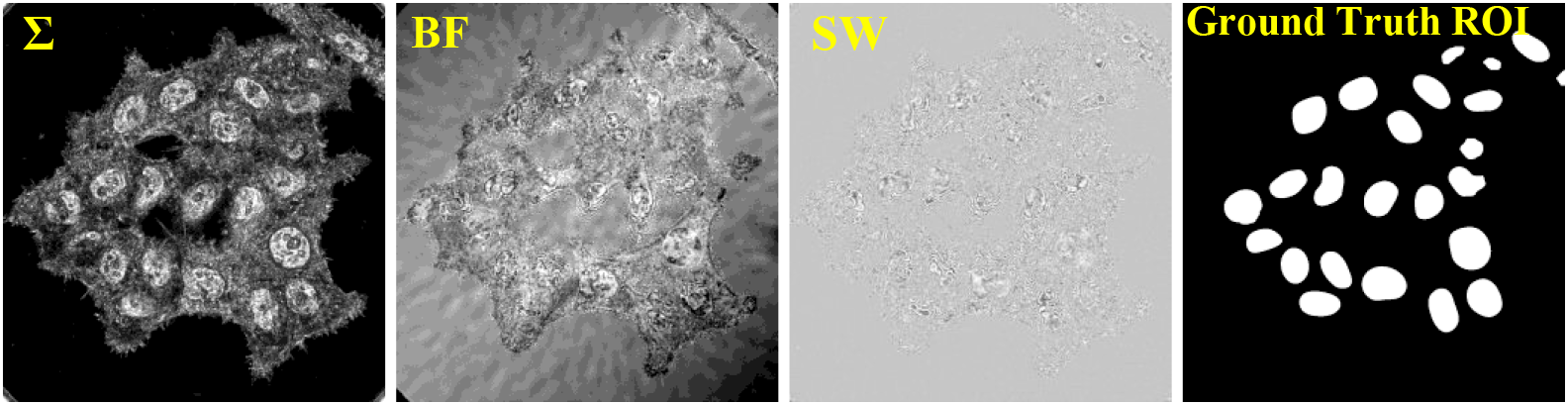
Sample training data. Three different csPWS feature image sets Σ (interference spectra) image, BF (brightfield) image, and SW (single wavelength) reflectance image, and their Ground Truth ROI mask.

### 2.6 Deep Learning Architecture

DL has revolutionized image segmentation, a crucial process in classifying each pixel of an image based on attributes such as color, intensity, or texture. The advent of Convolutional Neural Networks (CNNs) [34] and their capability to autonomously extract features from images through layers of convolution, pooling, and dense computations has set a new standard for image segmentation. Introduced in 2014, the Fully Convolutional Network (FCN) [35] marked a significant advancement by enabling end-to-end segmentation in a single forward pass. However, its fixed receptive field size limited its effectiveness across varying scales [36]. The U-Net architecture, which features a distinctive U-shaped design, was developed to address these limitations, particularly for complex biomedical image segmentation tasks [19]. U-Net is very efficient for complex medical image segmentation due to its ability to handle variations in image detail and scale, enhancing end-to-end segmentation capability. It uses multiple filters at various layers to extract and learn patterns from images, such as edges, textures, and shapes. However, standard U-Net struggles with capturing long-range dependencies in complex image datasets that require precise localization and contextual understanding for tasks such as semantic segmentation [37]. Its fixed feature aggregation and repeated pooling can blur finer details [38], affecting accuracy in segmenting the intricate structures of complex images. Attention mechanisms [39] have recently been introduced to overcome these limitations.

Attention mechanisms enhance U-Net by focusing on salient features that are critical for accurate segmentation while filtering out irrelevant and redundant data. This recalibration allows the network to better manage global and localized information, making it more effective in segmenting various objects within an image [40]. Leveraging this advantage, csPWS-seg uses a gated channel-based attention mechanism integrated into U-Net [20] for robust segmentation performance. This mechanism can control the activation of different channels based on the input that allows the model to dynamically adjust which features to emphasize, which is crucial in medical images where specific characteristics or shapes of the structures must be accurately identified and segmented. By selectively processing only the most relevant channels of information, gated mechanisms improve the model’s efficiency and accuracy, reducing the likelihood of misclassification in areas with subtle differences in texture or shape. Additionally, the adaptive feature selection capability of gated mechanisms helps in dealing with the variability in medical imaging caused by different imaging modalities or patient-specific factors [20]. This leads to more robust and reliable segmentation, which is essential for effective diagnosis and treatment planning in clinical settings.

The GCT-U-Net, which is the core of csPWS-seg, is shown in Fig. 2(a). It takes input 512×512 pixel size grayscale csPWS image and produces the segmented nucleus ROIs. U-Net’s core structure in GCT-U-Net is characterized by a contracting path that captures contextual information and a symmetrically expanding path that aids in precise localization. The contracting path, i.e., encoder - ConV1 to ConV4 block, each consists of two 3×3 convolutional layers having the same feature map size before and after the convolution operation followed by a 2×2 max pooling for reducing the network complexity and compressing the features. This encoder doubles the feature maps (16 to 128) and halves the image dimensions (512×512 to 64×64) as it moves deeper from one ConV block to the next, starting from ConV1 to ConV4, thus compressing spatial dimensions and capturing deeper contextual data from the input image. In the expanding path, i.e., the decoder - ConV6 to ConV9 block, each consists of a transposed convolution (or up-convolution) layers with twice the image dimension and halves the feature map followed by two regular convolutional layers through skip-connection with corresponding feature maps from the contracting path, facilitating precise delineation of the regions of interest by leveraging both high-level and local information. The deepest part of the network, ConV5, is designed to retain high-level features without further reducing spatial dimensions by using a max-pooling layer. This strategy preserves detailed spatial information crucial for accurately reconstructing image details during the upsampling process. At the end of the decoding process, we employed a 1×1 convolutional layer to produce the final segmented image. The proposed network has 23 convolutional layers. We also used dropout [41] layers within this network to prevent overfitting. The dropout rates were set at 0.1 for early layers and 0.2 and 0.3 for deeper layers. The lower dropout rates in initial layers preserve more information, while higher rates in deeper layers enforce stronger regularization to manage increased model complexity.

**Fig. 2.**
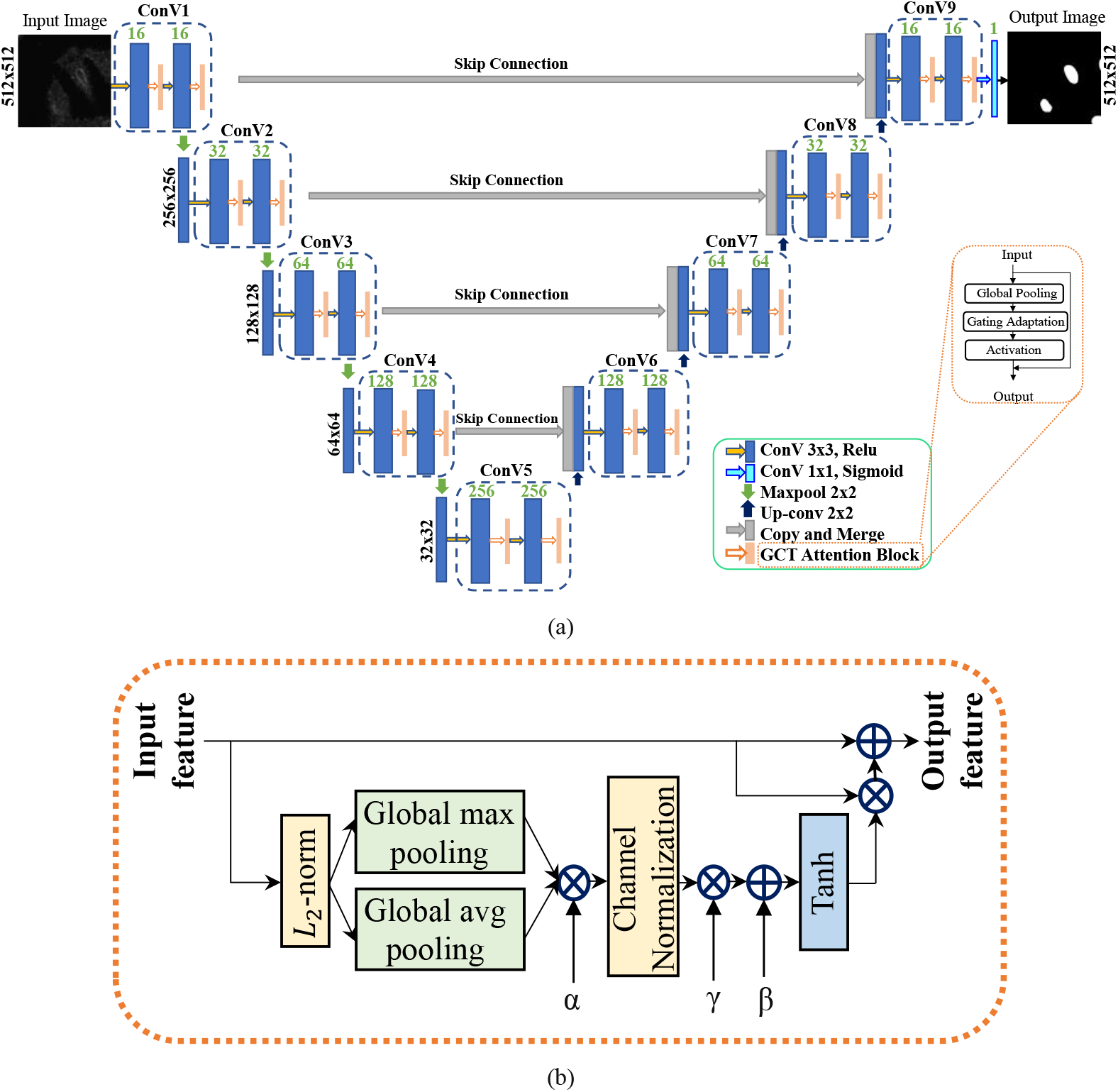
(a) Architecture of the proposed attention-based GCT-U-Net model used in csPWS-seg (b) Layer description of GCT attention Block.

The GCT (Gated Channel Transformation) blocks are integrated into both the contracting and expanding paths after each convolutional layer in each ConV block. This allows the model to refine features early, making downsampling and upsampling more effective. This ensures that the downsampled and upsampled features are fine-tuned using the recalibrated channel-wise features, enhancing the precision of the segmentation by adjusting the importance, or “weight,” of each feature channel in the convolutional feature maps. It does this by analyzing the overall information contained in each channel and deciding how much to emphasize or suppress that channel’s features. This decision-making process helps the network focus on more relevant features and suppress less useful ones, allowing for sharper and more accurate segmentations. The general GCT attention block is shown in Fig. 2(a), and its detailed version is in Fig. 2(b).

The trainable parameters α, β, and γ in the GCT block are responsible for additional feature transformation from L2 normalized output. The parameter, α, controlled the aggregation of L2-norm [42] (magnitude of feature vectors across spatial dimensions within a channel) global contextual information obtained via Global max and avg pooling operation. The channel normalization layer, then dividing the original feature vector by its L2-norm, scaling the features down to a unit vector, and preventing the dominance of particular features solely based on their scale rather than their relevance to the task to ensure no single channel dominates, which improves the generalization of the network. The gating mechanism adopted by parameters β and γ learns to ’open’ or ’close’ gates for certain information paths, effectively controlling the flow of feature-making emphasized or suppressed, optimizing the network’s responsiveness to relevant details while filtering out noise through the activation function. β functions as a bias in the gating mechanism, altering the significance threshold for channel features, thus shifting activation levels. γ, the gating weight, determines the impact of normalized channel values on gating, enabling dynamic adjustments in feature flow throughout the network. Thus, these parameters control the network’s ability to focus on more appropriate features while suppressing less useful ones. A Tanh activation function is then applied to the γ-weighted input. Finally, the results are combined to produce the output tensor. This approach not only enhances feature representation but also ensures that the most critical features have been extracted. This attention mechanism helps to overcome the intrinsic challenges of label-free csPWS data by enhancing feature extraction and improving the precision of segmentation of cell nuclei.

We also compare our csPWS-seg performance with the baseline U-Net and another attention-based SE-U-net model [26]. The detailed network architectures of both of these models are presented in supplementary section 1 (Fig. S1 and Fig. S2).

#### 2.6.1 Deep Learning Training Strategy and Loss Functions

For the training of all models, we first normalized the raw csPWS images using the min-max normalization, which transforms all pixels of the images into a decimal between 0 and 1. The goal of normalization is to make every data point have the same scale, so each feature is equally important. We then split the normalized csPWS data into the ratio of 90:10 for training and validation. The proposed model was trained and tested in Python using the TensorFlow 2.10 (Keras in the backend) DL framework. We use ReLU (Rectified Linear Unit) activation function [43]. The model was trained using ADAM optimizer [44] with the initial learning rate set to 1 × 10^−4^, with β1 and β2 parameters set to 0.1 and 0.001, respectively. This smaller learning rate results in slower training progression but provides good stability.

For the attention mechanism, the initial values of α, β, and γ were chosen in a way that the GCT can effectively integrate into the existing network without disrupting the stable training process. The authors in [18] suggest using the initial value of α as 1, which will preserve the original distribution of features across channels, making the training phase more stable. The γ and β were both initialized to 0. This setting, claimed by the author [18], will help in integrating the GCT into existing networks without immediately altering their behavior, providing a neutral starting point that will be updated through training.

All necessary packages and libraries were installed in the Windows operating system with Intel(R) Xeon (R) CPU @4.10GHz, 128GB RAM, and NVIDIA RTX A4500 to train and test the model to find the ROI, automatically calculate the number of the nuclei and draw the bounding box in each nucleus. Training of the csPWS-seg model with the combined dataset model took approximately 6 hours for 100 epochs with batch size 8. The training time in SE-U-Net and baseline U-Net was 5 and 4.5 hours, respectively, for the same epochs and batch size. Testing, on the other hand, was considerably faster, with each model taking an average of less than 10 seconds to segment all the nuclei within the image of size 1024 ×1024 pixels. These times highlight the efficiency of our model in operational settings.

To compare the optimal training and testing performance of our model, we employed distributions-based (binary cross-entropy (BCE) and focal) and region-based (dice and Jaccard) loss functions tailored for segmentation tasks. The BCE is a commonly used loss function for binary classification tasks. It measures the difference between two probability distributions - the predicted probabilities and the actual binary outcomes (labels). The BCE loss function is mathematically defined as:

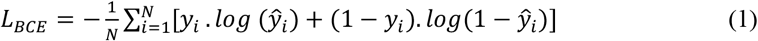

where *y*_*i*_ is the true label and 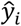 is the predicted probability for the *i*-th pixel.

Similarly, focal loss is an adaptive loss function designed to address the class imbalance by focusing on hard-to-classify examples (smaller objects) and down-weights well-classified examples, i.e., background. It modifies the standard cross-entropy loss to reduce the relative loss for well-classified examples, putting more emphasis on correcting misclassified data, resulting in a dynamic scaling of the cross-entropy loss, where the scaling factor decays to zero as confidence in the correct class increases, making it more robust to easy negatives that dominate large datasets. This mechanism helps to enhance the learning focus towards hard, misclassified examples, thus improving the performance of the model on more complex and varied datasets. The focal loss function is mathematically defined as:

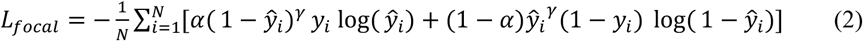

where *γ* is the focusing parameter and *α* is the balancing factor which helps in prioritizing harder-to-classify examples and balances the class distribution. This is especially beneficial in scenarios with imbalanced datasets. For our training, the values of *α* and *γ* are set to 0.25 and 2, respectively.

The dice loss is another loss function often used in segmentation tasks. This measures the overlap between the prediction and the ground truth, where a perfect overlap results in a Dice Similarity Coefficient (DSC) of 1 and no overlapping results in 0. As a loss function, it is formulated as (1 − DSC), transforming it into a minimization problem where the objective is to reduce the loss, thus maximizing the DSC. The DSC is expressed mathematically as:

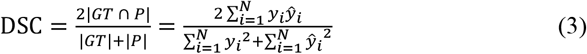

where *GT* represents the ground-truth and *P* represents the predicted images. Both forms of the formula represent the DSC, yet their applications may vary based on whether the model outputs are probabilistic or binary and whether the context is model training or performance evaluation. Typically, during model training, the output may not be strictly binary, which necessitates a modified approach to the DSC to ensure effective gradient-based optimization. This adaptation allows for continuous updating of optimization parameters in the model even when predictions are probabilistic. The dice loss formula, therefore, can be defined as:

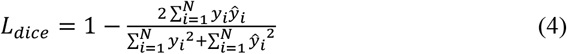

This formulation ensures that as the dice loss decreases, the overlap between the predicted segmentation and the ground truth increases, indicating improved model performance.

Similar to dice loss, the Jaccard loss is another common loss function for evaluating the performance of segmentation models, particularly useful for assessing how well the predicted segmentation matches the actual labels results in a Jaccard coefficient, also referred to as Intersection over Union (IOU). When the Jaccard coefficient approaches 1, within a value range of 0 to 1, it indicates that the prediction closely aligns with the ground truth. As a loss function, it is formulated as (1 − Jaccard coefficient). The Jaccard coefficient and the Jaccard loss can be expressed as:

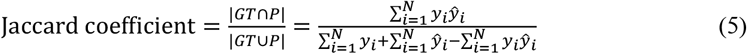

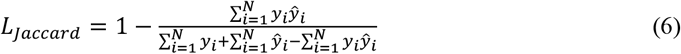

#### 2.6.2 Evaluation Metrics

The segmentation performance of the trained model was evaluated using two commonly used evaluation metrics: IoU (Intersection over Union) and Dice Similarity Coefficient (DSC), also called the F-1 score for semantic segmentation. The range of these values is between 0 to 1.

The higher the value, the better the model performance. The IoU score provides a quantitative measure of the percentage overlap between the predicted region of interest (ROI) and the ground truth, effectively indicating how much the predicted segmentation covers the actual area of interest. On the other hand, the DSC score points towards its effectiveness in balancing precision and recall for segmentation tasks, combining both to assess the accuracy of the segmented regions while penalizing both false positives and false negatives. The DSC score is a robust indicator of the model’s overall performance in distinguishing and correctly classifying each pixel of the target structure. The IoU and DSC can be defined as:

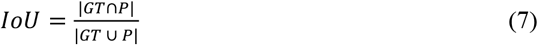

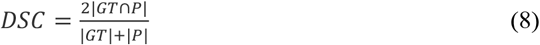

where *GT* represents the ground-truth ROI and *P* represents the predicted ROI of test csPWS images. Also, the |*GT* ⋂ *P*| and |*GT* ⋃ *P*| represents the common elements and total number of unique elements, respectively, between the sets of *GT* and *P*. The visualization of [Eq. 7] of IoU is shown in Fig. 3.

**Fig. 3.**
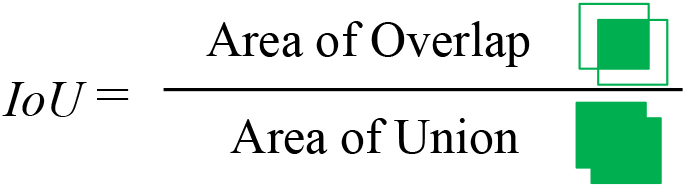
Visualization of IoU

## 3. Results and Discussion

### 3.1 Performance evaluation of csPWS-seg trained on various csPWS feature image data

We evaluated the csPWS-seg nuclei segmentation performance of live-cell HCT 116 imaging data on four different trained models. The four models include the model trained on BF image only, Σ image only, SW image only, and the collection of BF, Σ, and SW images. More specifically, first, we trained the csPWS-seg using the training data with individual feature images only. In this case, we trained three separate models using BF, Σ, and SW images-only training datasets. In this case, no feature images were mixed during training. These individually trained models are then tested with new test feature images from all. We then recorded evaluation scores: IoU and DSC scores, as shown in the box plot of Fig. 4 (a) and (b). The box plot includes the distribution of these scores from 18 (6 from each feature image) different randomly selected test images. Among individually trained models, the median IoU and DSC score of the SW-only trained model is the highest and shows satisfactory performance when tested with data from any feature images. Additionally, the higher IoU values on the box plot for each modality are from the test images from the same group. For example, the Σ-only model, when tested with Σ images, will have higher IoU and DSC scores with better segmentation results. When this model is tested with images from different groups, such as BF, the IoU and DSC scores are usually low. This is because the neural networks learn features from the same class of images and fail to provide satisfactory results when tested with images from different groups having different features. Nevertheless, the SW-only trained images provided satisfactory results on all kinds of test feature images due to their unique characteristics, as explained in section 2.5. The segmentation results for individually evaluated models are shown in Fig. S3 of the supplementary section 2.1.

**Fig. 4.**
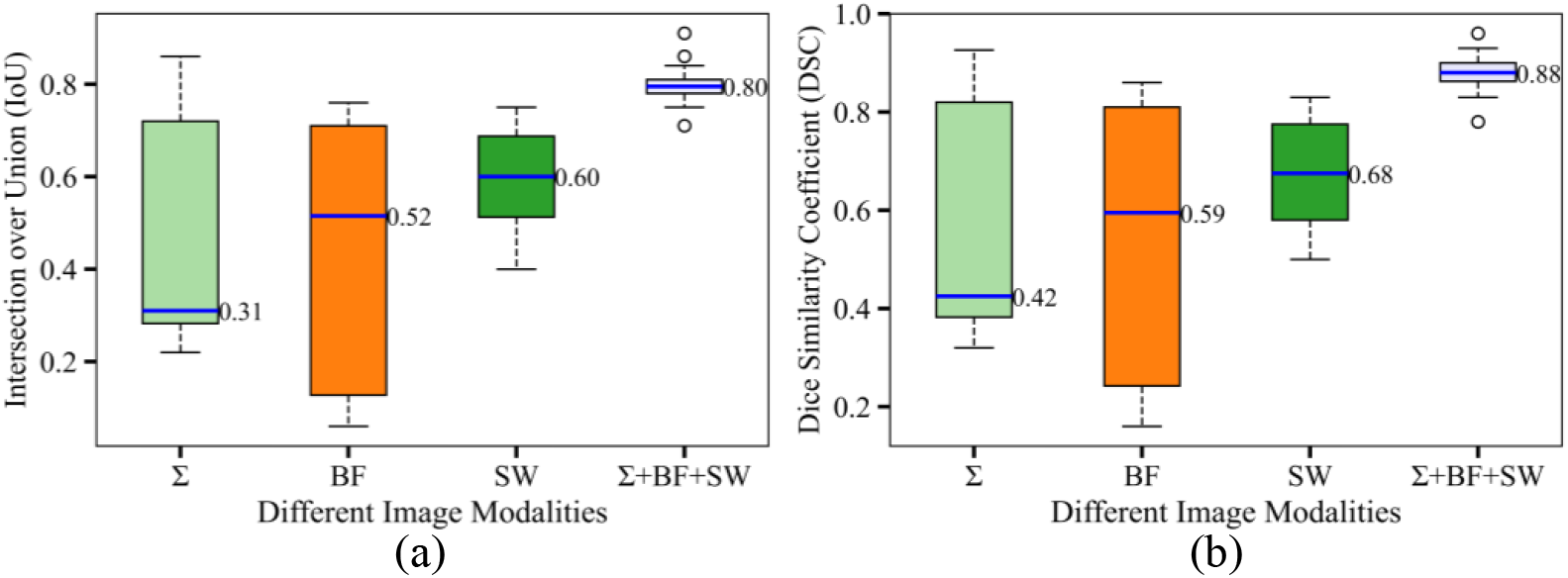
(a) IoU and (b) DSC score of csPWS-seg trained on various csPWS feature image datasets.

To improve the performance of the individually trained model, we trained a new model mixing images from all three feature images, i.e., the collection of BF, Σ, and SW images. We then tested this model with any random csPWS feature image and performed the performance evaluation. The median IoU and DSC-score for this trained model improved significantly and are 0.80 and 0.88, respectively (Σ+BF+SW box plot in Fig. 4 (a) and (b)), showing excellent segmentation performance on any group of feature imaging data. Notably, SW images, when tested on combined trained models, have the highest true positive (TP) values (Table S1 of the supplementary section 2.5), showing the advantage of SW feature images for csPWS nuclei segmentation analysis.

Comparative visual analysis results of sample test images on the model trained in combined datasets are illustrated in Fig. 5, where (a) shows the input test images, and (b) shows the test images superimposed with the prediction segmented nuclei ROI (red) with boundaries (green). The IoU and DSC values of the segmented nuclei are also presented. The zoomed-in views of a randomly selected nucleus in Fig. 5 (c) emphasize the textural and structural details of the nucleus along with accurate prediction, visualizing highly accurate segmentation using csPWS-seg. The green contours in the zoomed predicted ROIs delineate the precision of segmentation with an accurate boundary region. This visualization highlights the csPWS-seg superior nuclei segmentation performance on complex csPWS feature imaging data, automating the accurate nuclei segmentation task for chromatin analysis.

**Fig. 5.**
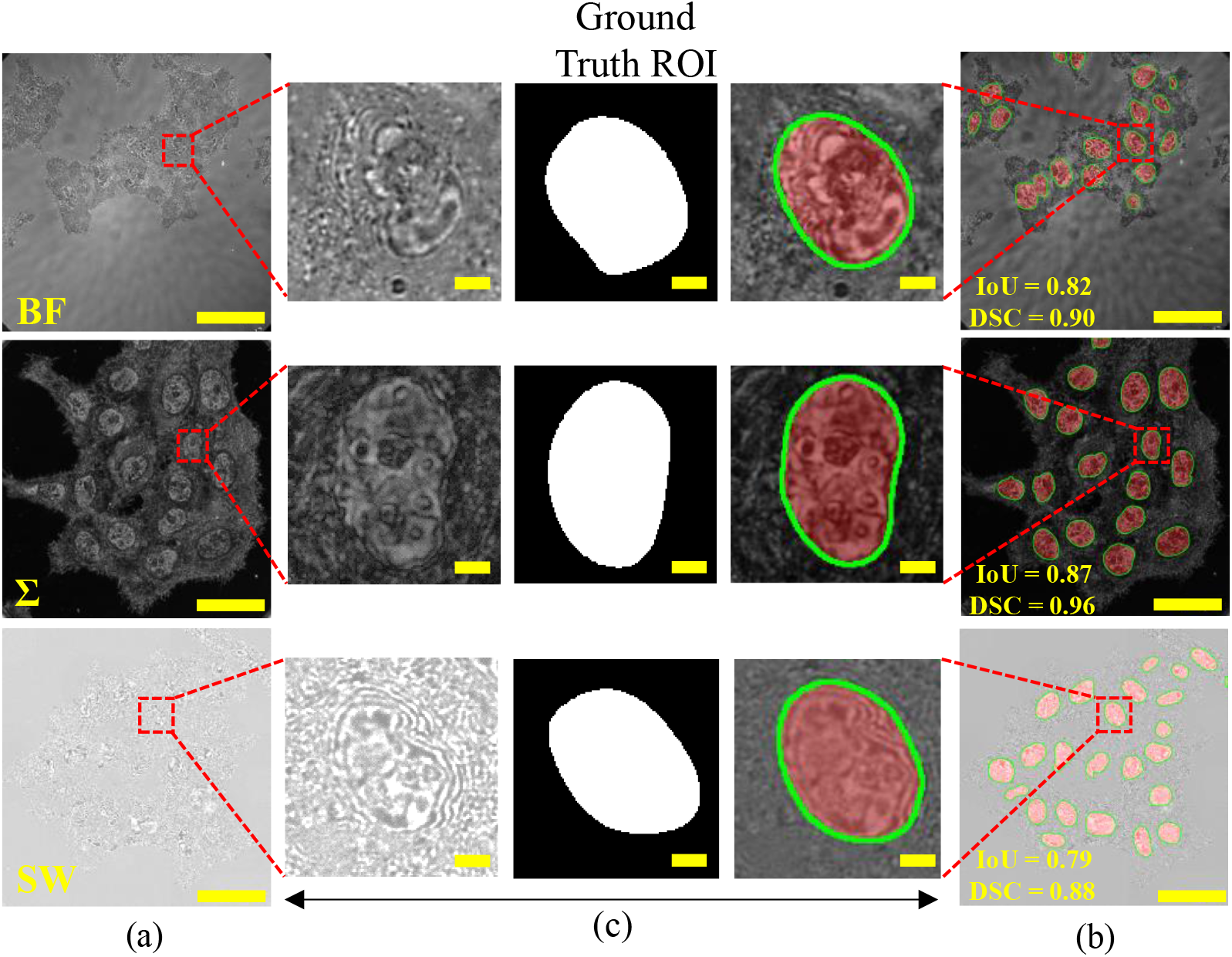
csPWS-seg performance analysis. (a) Test images. (b) Test image superimposed with the predicted ROIs. (c) Zoom in view of selected nuclei from (a) and (b). Ground truth ROI is in the middle. The predicted ROIs are highlighted in red, and boundary regions are shown in green. Scale bar: 50μm for test image and 25μm for zoom-in image.

We then compared the csPWS-seg performance with the baseline U-Net model and another popular attention-based model, SE-U-Net. The comparative performance analysis results of all three DL models are illustrated in Fig. 6. We can observe that the attention-based model, including the csPWS-seg model, consistently showed improved median IoU and DSC scores over the baseline U-Net, indicating enhanced segmentation capabilities offered by their attention mechanisms. Some of the gradual improvement in predicted ROIs, as well as the boundary regions over the model, has been indicated by the white arrows. These results indicate that the csPWS-seg leveraging the great benefit of the attention mechanism offered significant improvement in segmentation accuracy. Further analysis of these three DL models with different loss functions is presented in section 3.3.

**Fig. 6.**
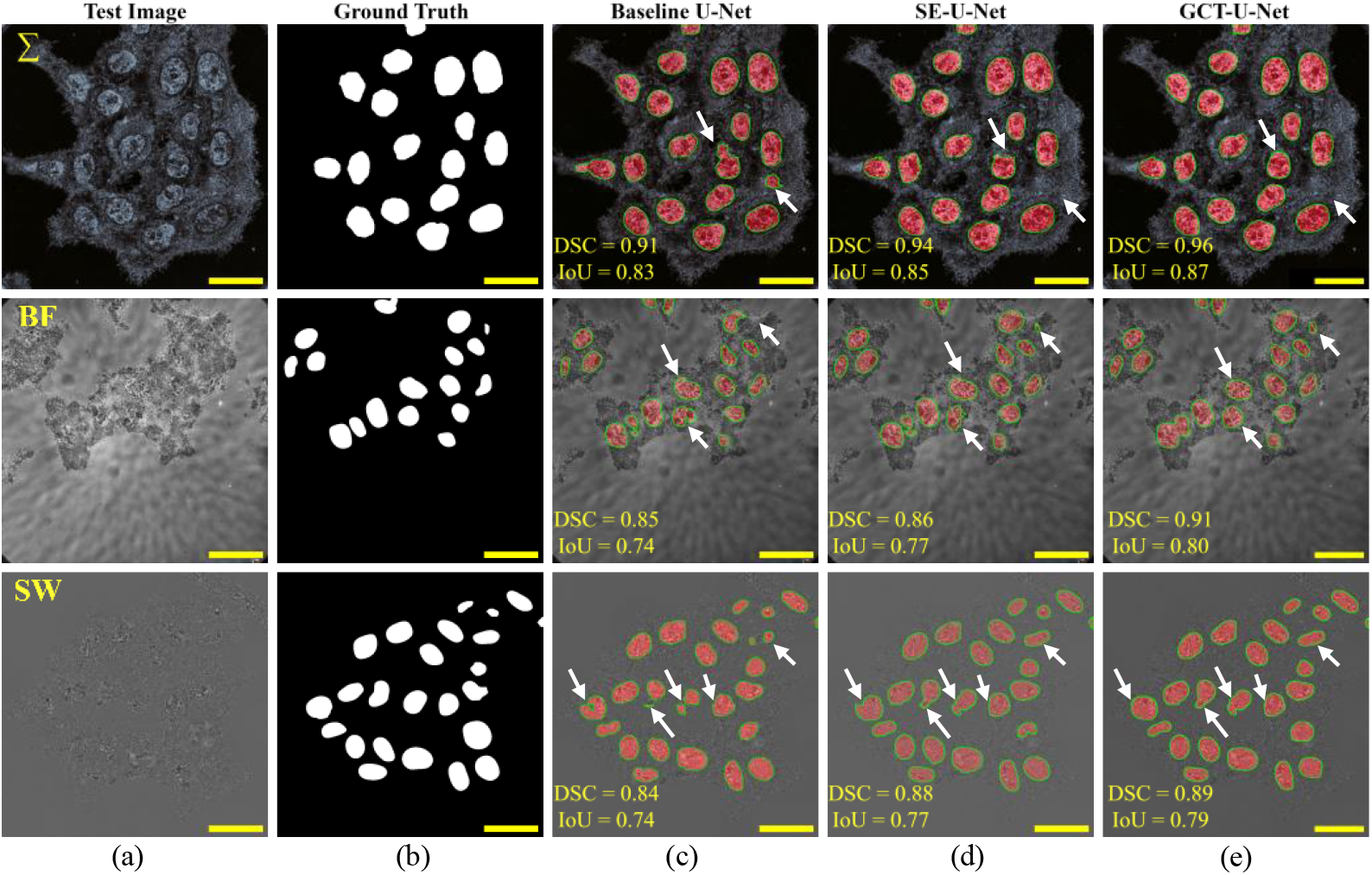
Performance analysis on three different DL models. (a) Test images. (b) Ground-truth ROIs. (c)-(e) Predicted ROIs superimposed with the test images for the baseline U-Net model, SE-U-Net model, and csPWS-seg, respectively. The predicted ROIs are highlighted in red, and boundary regions are shown in green. Scale bar: 25 μm

### 3.2 Performance evaluation on different loss functions

The loss function plays a pivotal role in the training of DL models for image segmentation. We analyzed the performance of our DL models over several popular segmentation loss functions with the goal of obtaining the best segmentation loss function suitable for csPWS microscopy image nuclei segmentation. For that, we chose distribution-based (BCE loss and focal loss) and region-based (dice-loss and Jaccard loss) loss functions and trained our DL models separately by using one loss function at a time.

The training and validation loss curve of the distribution-based optimization loss function for csPWS-seg for the collection of BF, Σ, and SW image datasets are shown in Fig. 7 (a) and (b). Figure 7 (a) shows the loss curve for the focal loss function, and Fig. 7 (b) shows the loss curves for the BCE loss function. It can be seen that csPWS-seg shows a significant decrease in train and validation loss with respect to a number of epochs for both focal loss and BCE loss. It shows a good convergence pattern compared to other individual feature images trained models (as shown in Fig. S4 and Fig. S5 in supplement sub-section 2.2 of section 2), indicating that incorporating attention mechanisms enhances its feature discrimination capabilities. In both graphs, the train and validation loss decreased gradually up to about 70 epochs and then plateaued until 100 epochs. This continuous and steady decline in loss without significant fluctuations suggests that the model is not overfitting and is generalizing well to new data and provides a great advantage in segmenting the cell nuclei. Therefore, it can be concluded that the proposed csPWS-seg training strategy and loss function get the optimal performance. Additionally, the training and validation curves for the distribution-based loss function for individually trained feature image models are presented in Fig. S6 of the supplementary section 2.3.

**Fig. 7.**
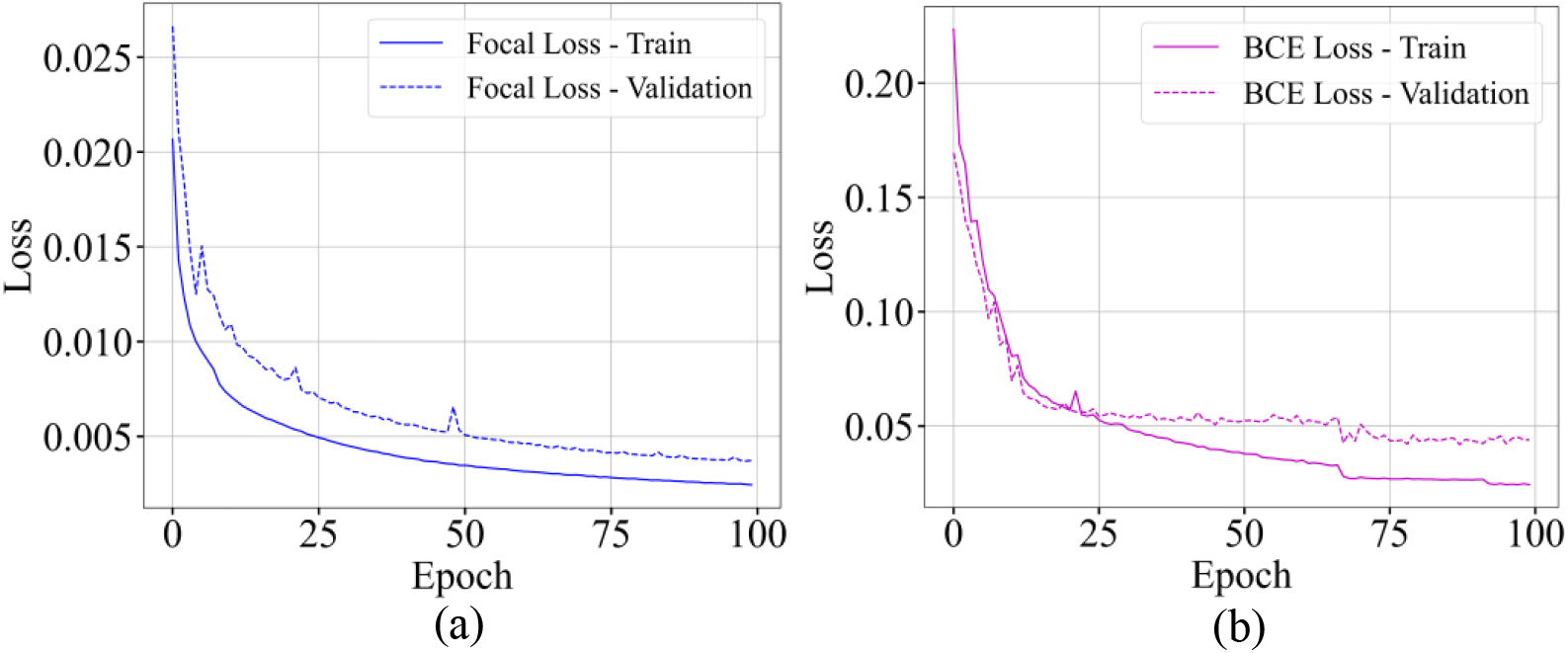
Train and validation loss curve on (a) focal loss and (b) BCE loss on csPWS-seg during training.

Similarly, the training and validation loss curve for the region-based loss functions for the collection of BF, Σ, and SW image datasets are shown in Fig. 8. Figure 8 (a) shows the loss curve for dice loss, and Fig. 8 (b) shows the loss curves for Jaccard index. Although the loss value is decreasing, it is relatively higher compared to the distribution-based loss functions of Fig. 7. Also, some noticeable abrupt drops at around 20 epochs in loss values can be observed in Fig. 8 (a) and (b). The reason for this is that the model reaches a point where predictions start to overlap significantly with the ground truth ROIs, where both loss performances improve noticeably. It may also be because of wide variations in the distribution of the training sample (such as variations of the wide range of features in the samples in Σ and SW) that contribute to noticeable changes in their objective function. Thus, when the model calculates the gradient from the features on relatively easy (Σ) to hard (SW) samples, it can cause sudden shifts in the loss function. Additionally, the training and validation curves for the region-based loss function for individually trained feature image models are presented in Fig. S7 of the supplementary section 2.4.

**Fig. 8.**
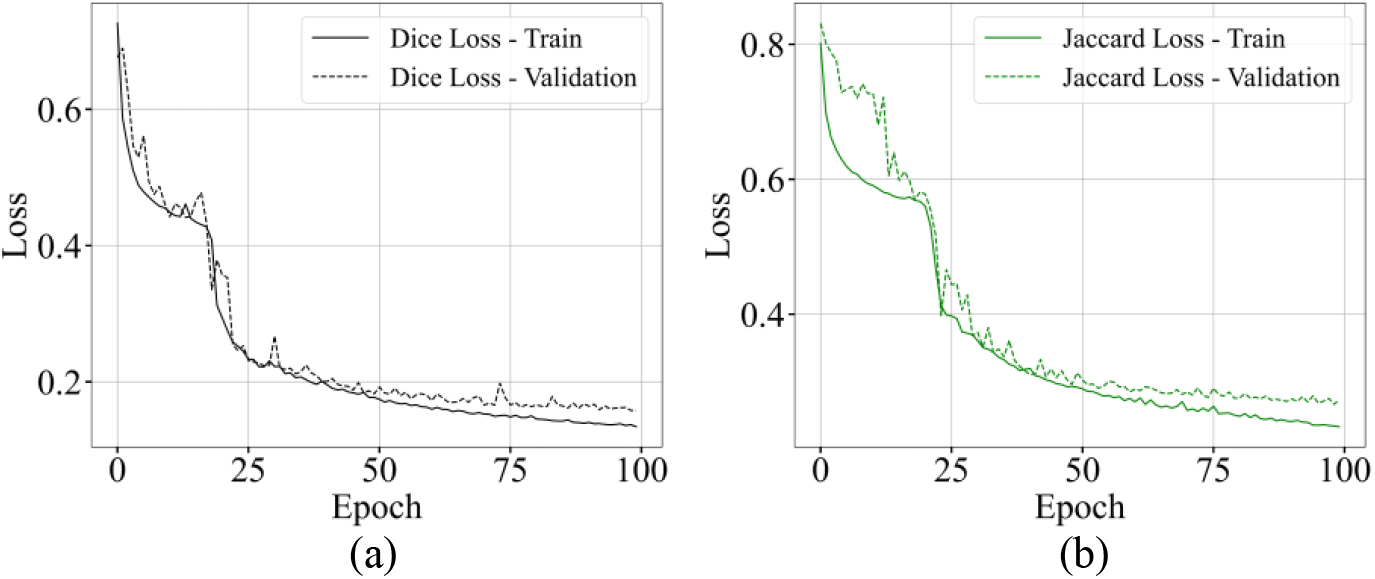
Train and validation loss curve on (a) dice loss and (b) Jaccard loss in csPWS-seg during training.

The comparative study with both distribution and region-based loss functions shows that the loss value for focal loss is lowest indicating that the csPWS-seg is learning much more effectively using this loss compared to others, providing the highly reliable segmentation of nuclei. Thus, the results presented in all Figs. 4-6 are from the model trained with the focal loss function.

### 3.3 Performance comparison of different DL architectures on different loss functions

We evaluate and compare the performance of distribution-based and region-based loss functions with all three DL architectures (U-Net, SE-U-Net, and GCT-U-Net). The box plot showing the IoU and DSC scores for the three DL architectures trained separately on four different loss functions is shown in Figs. 9 and 10 (a-d)
. These scores are from the models trained with combined BF, Σ, and SW images. The median evaluation scores are indicated in each box plot. The results show that SE-U-Net and GCT-U-Net consistently show improved IoU and DSC scores over the baseline U-Net, indicating enhanced segmentation capabilities due to their attention mechanisms and appropriate segmentation loss function. Overall, GCT-U-Net trained with focal loss performs best, with the highest IoU score of 0.80 and DSC score of 0.89. Further, training and validation performance curves for these loss functions with individually trained feature images for all three DL architectures are presented in Figs. S4 and S5 and explained in section 2.2 of supplementary 1.

**Fig. 9.**
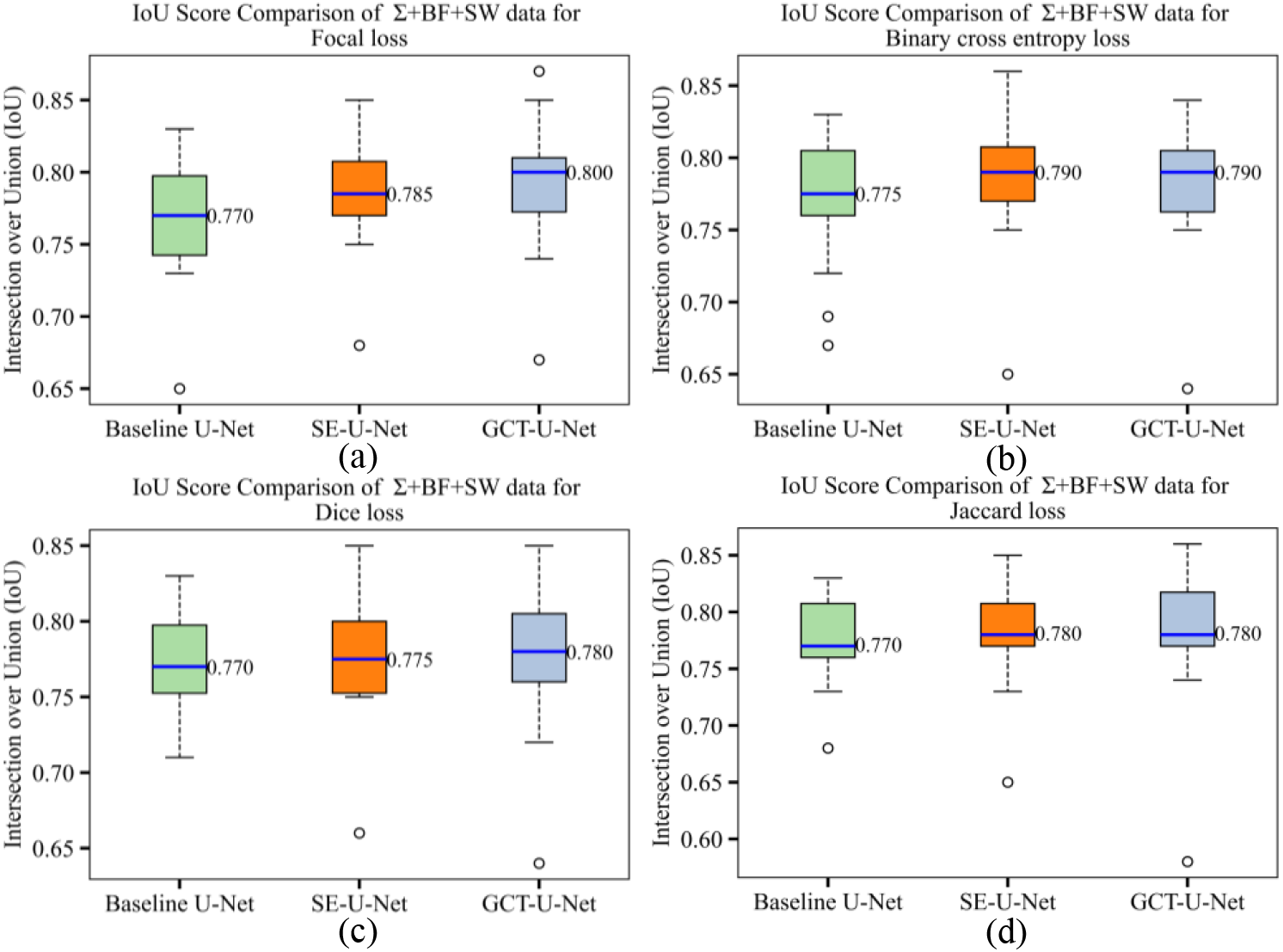
Box plots comparing the performance of different DL architectures using IoU metric for (a) Focal loss, (b) BCE loss, (c) dice loss, and (d) Jaccard loss functions.

**Fig. 10.**
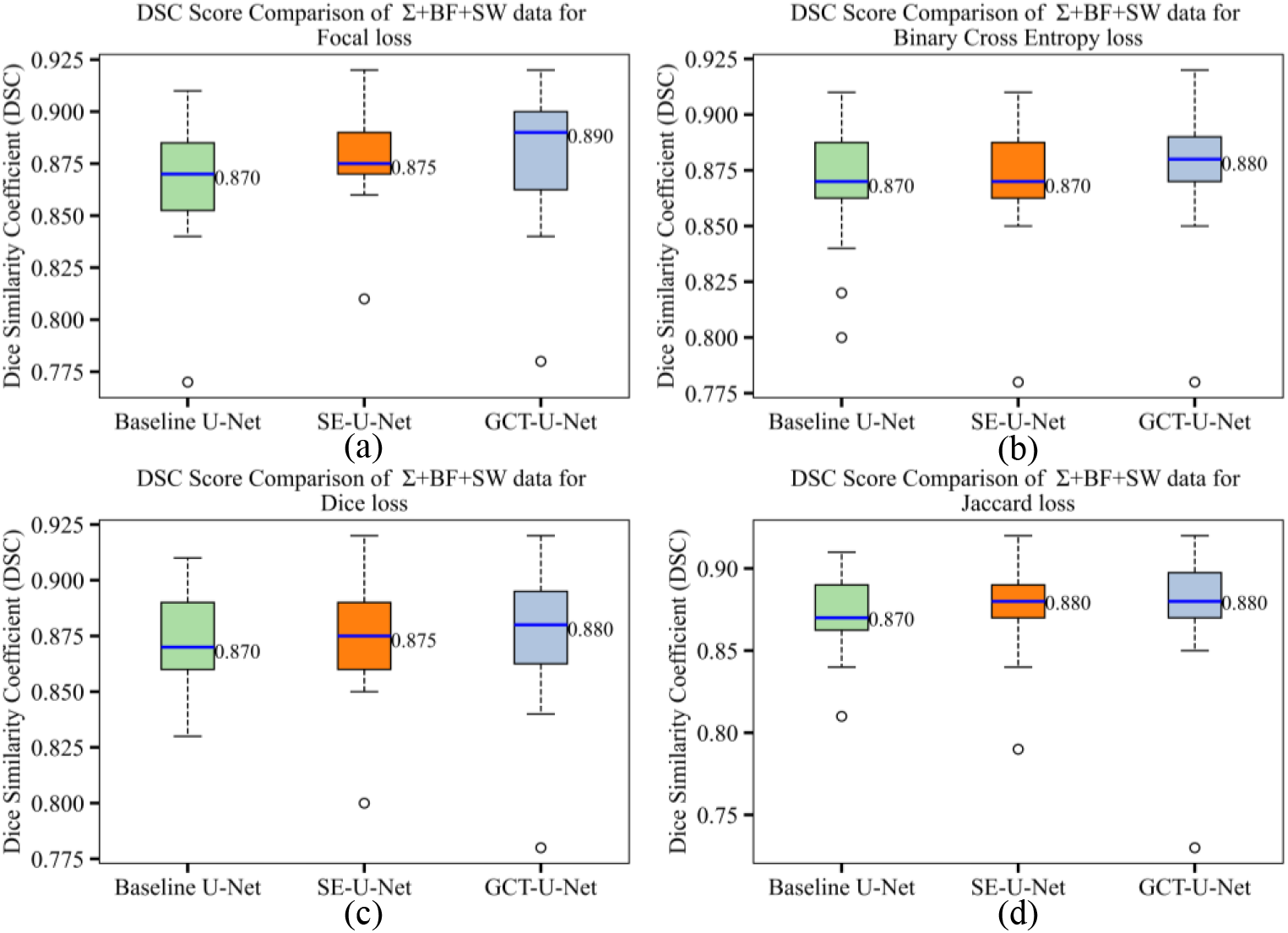
Box plots comparing the performance of different DL architectures using DSC metric for (a) focal loss, (b) BCE loss, (c) dice loss, and (d) Jaccard loss functions.

## 4. Conclusions

In this work, we propose a novel DL technique termed csPWS-seg to automate and accelerate the accurate nuclei segmentation for live cell csPWS microscopy imaging data. csPWS-seg leverages the combination of gated attention mechanisms and focal loss functions integrated with a baseline U-Net architecture, providing a superior segmentation task. csPWS-seg outperformed the baseline U-Net and SE-U-Net in terms of segmentation accuracy when evaluated using the HCT116 human colon cancer cell line data. The csPWS-seg performance was evaluated by using metrics IoU and DSC scores, achieving a median IoU of 0.80 and a DSC score of 0.88. We also compared the performance with several loss functions and found that focal loss notably enhanced the segmentation performance. This is because the focal loss function effectively prioritized the segmentation of challenging regions by adjusting the emphasis on hard-to-classify examples. csPWS-seg offers approximately 60-fold faster nuclei segmentation, avoiding error-prone and time-consuming manual segmentation. In addition, csPWS users can use any of the BF, Σ, or SW images to segment nuclei. Our approach will greatly ease and expedite the segmentation task for both novel and experienced csPWS users. We hope that automated, fast, and accurate protocols for the characterization of the nucleus will disseminate csPWS microscopy to the broad biomedical research community, including segmenting and analyzing other cell lines and tissues for clinical applications.

Although csPWS-seg demonstrates high effectiveness, it occasionally misidentifies non-nuclei regions smaller than typical nuclei, primarily due to unclear boundaries in actual cell images. This issue underscores the need for further model refinement, particularly due to the limited size of the training dataset and the inherent variability in biological samples. Additionally, the presence of noise and low signal-to-noise ratio due to label-free images complicates the segmentation process. Nevertheless, this study not only advances the technology for analyzing cellular structures but also contributes to the broader understanding of cell morphology and its implications in cancer diagnostics and treatment. We anticipate that continued enhancements to DL techniques and progress in csPWS instrumentation and imaging systems will further refine the accuracy of cell nuclei segmentation and analyses, thereby supporting more precise detection, diagnostics, and treatment strategies for cancer.

## Acknowledgments

The authors would like to thank Joshua Pritchard for helping with cell culture.

## Data availability

Code and data relevant to this article are not publicly available but may be obtained from the authors upon reasonable request.

## Supplementary document

See Supplementary 1 for supporting content.

