## Supplementary material for "Deep Learning-driven Automatic Nuclei Segmentation of Label-free Live Cell Chromatin-sensitive Partial Wave Spectroscopic Microscopy Imaging": csPWS-seg Supplimental File Final.pdf

### 1. Detailed Architecture of Baseline U-Net and SE-U-Net

#### Baseline U-Net Architecture

The U-Net [1] employs an encoder-decoder architecture that enhances spatial accuracy and ensures more precise segmentation outcomes. In the encoder, the image is progressively downscaled while increasing feature depth to extract high-level information. Conversely, the decoder upscales these features and reconstructs the segmentation map by gradually increasing spatial dimensions. U-Net's distinctive skip connections [2] bridge the encoder and decoder blocks, transferring spatial and low-level feature information lost during downsampling back to the corresponding decoder layers. This symmetrical structure, along with the integration of lost low-level details via skip connections, allows for efficient end-to-end segmentation, making it particularly effective in medical imaging contexts. The adopted baseline U-Net architecture for csPWS [3,4] image nuclei segmentation is shown in Fig. S1.

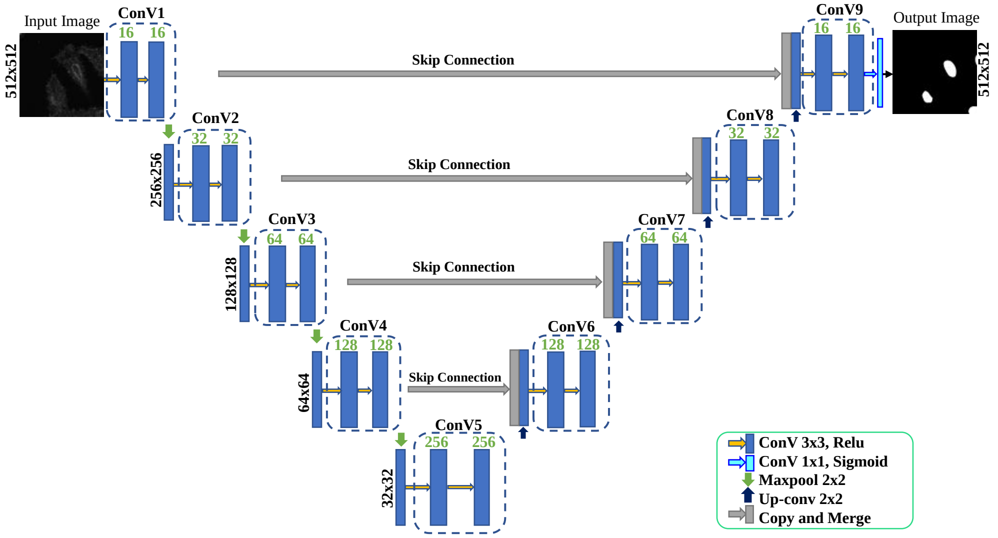


Fig. S1. Architecture of the baseline U-Net model.

The baseline U-Net architecture takes csPWS microscopy images sized 512×512 pixels, producing a nuclei segmentation map of the same size due to convolution operations. It utilizes 3×3 convolutional filters throughout its encoder-decoder structure (except the last decoder layer where we have used 1×1 to generate pixelwise segmented ROI), with 2 × 2 max pooling in the encoder to reduce dimensions and 2 × 2 up-convolutions in the decoder to restore them. The number of feature maps is doubled and the image size is halved than its previous ConV block in the encoder and a reverse process occurs in the decoder block. In Fig. S1, the number of image sizes and feature maps are shown at the left side and upper of each convolution layer, respectively. Also, each decoder block is concatenated with the result of the downsampling at corresponding spatial scales through a skip connection. Dropout layer [5] is added after each ConV block which helps to prevent overfitting by randomly deactivating a portion of neurons during training, ensuring the network learns robust features from our csPWS data. We used a dropout rate of 0.1 at the initial few layers of encoders - ConV1 & ConV2, and their corresponding decoder block. Meanwhile, in the deeper encoder and their corresponding decoder layers, a slightly higher dropout rate of 0.2 is used.

**SE-U-Net Architecture**

The SE-U-Net [6] integrates the Squeeze-and-Excitation (SE) [7] block in the baseline U-Net to incorporate the attention mechanism. The SE blocks in the convolutional neural network (CNN) enhance the network's ability to perform dynamic channel-wise feature recalibration. By using global average pooling, these blocks squeeze spatial dimensions to form channel descriptors, which are then processed through a gating mechanism involving fully connected layers. This process scales the original features based on their relevance, enhancing the network’s feature discrimination capabilities. The SE mechanism is shown in Fig. S2(a). By integrating these blocks into our baseline U-Net architecture, the network benefits from both spatial accuracy and enhanced feature representation. We placed this attention block at the end of each convolution layer. This clever arrangement leverages freshly processed features for immediate and effective enhancement, optimizing the network’s performance. The detailed model architecture, including the integration of these attention blocks, is illustrated in Fig. S2(b).

In the squeeze phase, SE blocks generate channel-wise statistics i.e. channel descriptor $z$, by extracting global information from each of the channels (during convolutional operations in the U-Net architecture, multiple filters at each layer generate various feature maps, or "channels,") of the input image by extracting information outside the receptive field of the convolution filter. It is computed as the average of all the feature maps in that channel. The excitation phase uses these statistics to recalibrate the input feature maps through a gating mechanism with the sigmoid activation function. The squeeze and excitation operations are given by [Eq. S1 and S2], respectively.

$$z_c = \frac{1}{H \times W} \sum_{i=1}^{H} \sum_{j=1}^{W} X_{i,j,C} \quad \text{(S1)}$$

$$s = \sigma\big(g(z, W)\big) = \sigma\big(W_2\, \delta(W_1, z)\big) \quad \text{(S2)}$$

where $H$, $W$, and $C$ denote height, width, and number of channels of input csPWS image, respectively. The excitation is done by reducing the dimensionality by a factor $r$ (reduction ratio, which controls the degree of dimensionality reduction in the gating mechanism), and then restoring it, followed by a sigmoid activation $\sigma$. In [Eq. S2], $g$ represents the gating mechanism, $\delta$ denotes the ReLU activation function, $W_1 \in \mathbb{R}^{\frac{C}{r} \times C}$ and $W_2 \in \mathbb{R}^{C \times \frac{C}{r}}$ are the weights of the two FC layers, and $s$ is the output scaling vector, which is then used to scale the original feature maps. This means each channel $C$ of the feature maps $X$ is multiplied by the corresponding scalar $s_c$ from the activation vector. This operation adjusts the intensity of the features in each channel based on the importance given by the SE block, yielding the recalibrated feature maps $X_c'$ that is given as:

$$X_c' = s_c \,. X_c \tag{S3}$$

The integration of the SE block with the baseline U-Net ensures the retention of critical channel-specific details that might otherwise be overlooked.

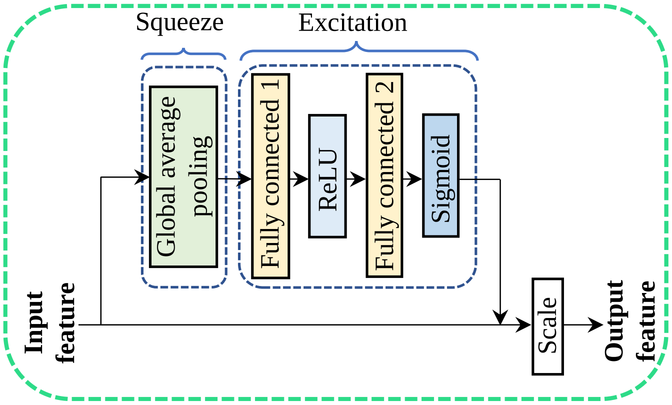


(a)

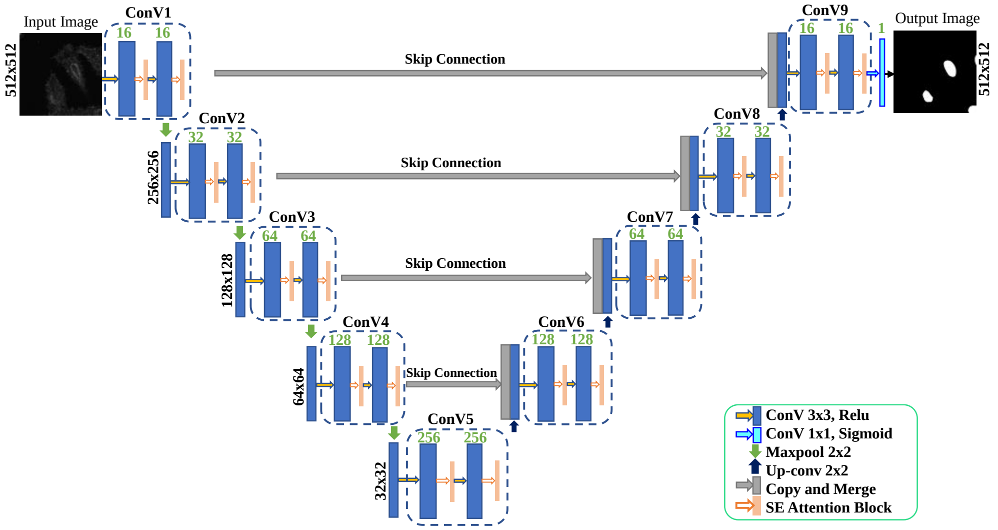


(b)

Fig. S2. (a) Layer description of SE attention Block. (b) The architecture of the attention-based SE-U-Net model.

### 2. Performance Evaluation

#### *2.1 Performance evaluation of csPWS-seg model on separately trained csPWS feature data*

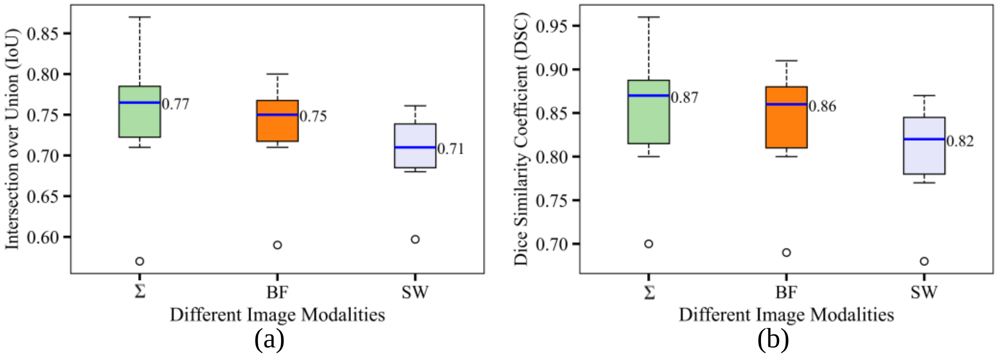


Fig. S3. Performance analysis of csPWS-seg trained over various csPWS feature images separately. (a) IoU and (b) DSC score.

Figure S3 illustrates the comparative performance of the csPWS-seg segmentation model when trained and tested on individual feature images – BF, Σ, and SW – using two key evaluation metrics: Intersection over Union (IoU) and Dice Similarity Coefficient (DSC). The box plots display the statistical distribution of these metrics, offering insights into the consistency and effectiveness of the segmentation process under varied feature images. Σ images show the highest median IoU (~ 0.77) and DSC (~ 0.87), indicating a robust segmentation capability. This superior performance is likely attributable to the detailed textural information inherent in Σ images, which aids in distinguishing complex cellular structures more effectively. Conversely, SW images exhibit the lowest median IoU (~ 0.71) and DSC (~ 0.82), indicating challenges in segmenting structures solely based on SW image data, where certain features may not be as obvious. These findings highlight the significant impact of feature image selection on the segmentation performance of csPWS imaging data. Combining different types of image features and mixing all of them for training improved the accuracies, as shown in the main text section 3.1.

*2.2 Training and validation performance with different loss functions for all adopted DL architectures*

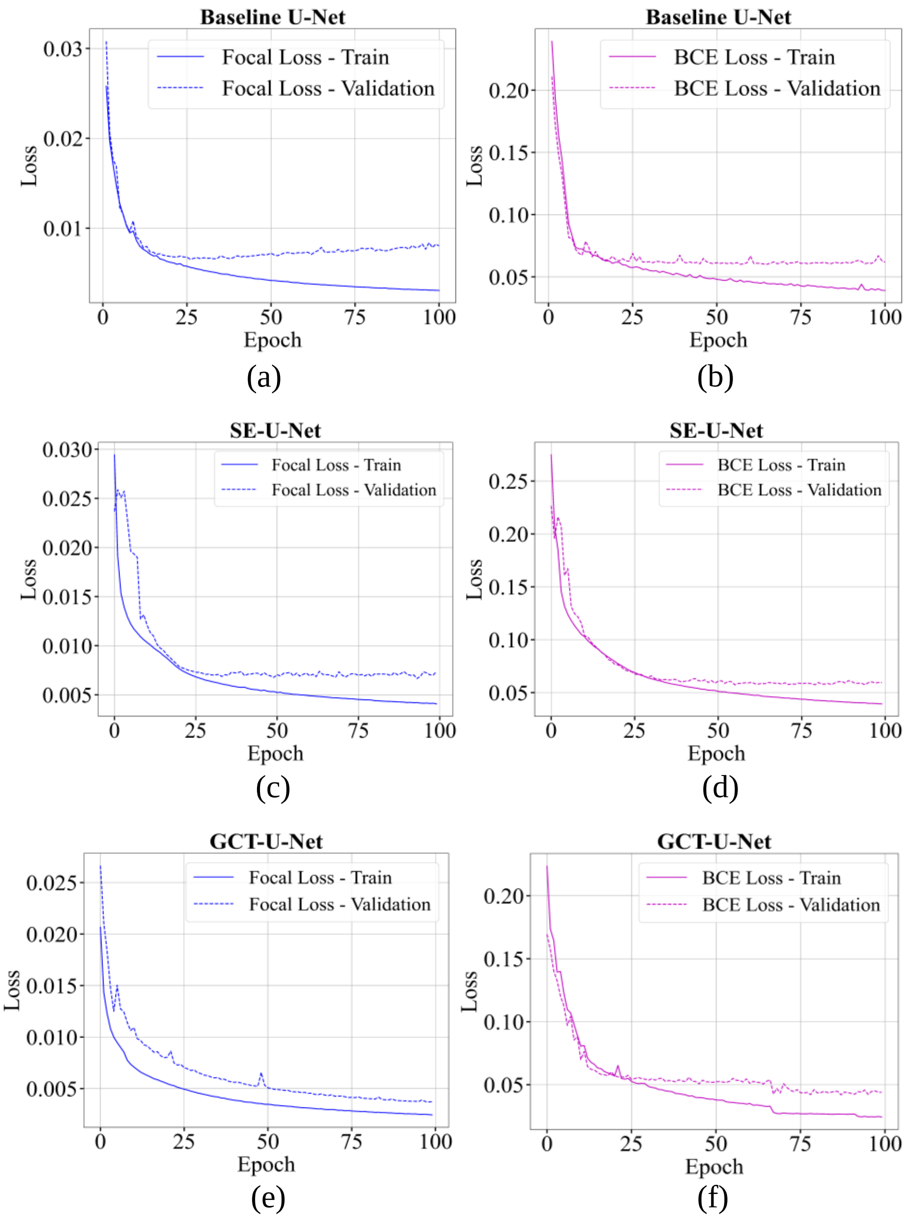


Fig. S4. Train and validation loss curve with respect to the number of training epochs on focal loss and binary cross entropy (BCE) loss for combined image modalities (Σ+BF+SW) for all DL architectures. **(a, b)** Baseline U-Net, **(c, d)** SE-U-Net, and **(e, f)** GCT-U-Net (csPWS-seg).

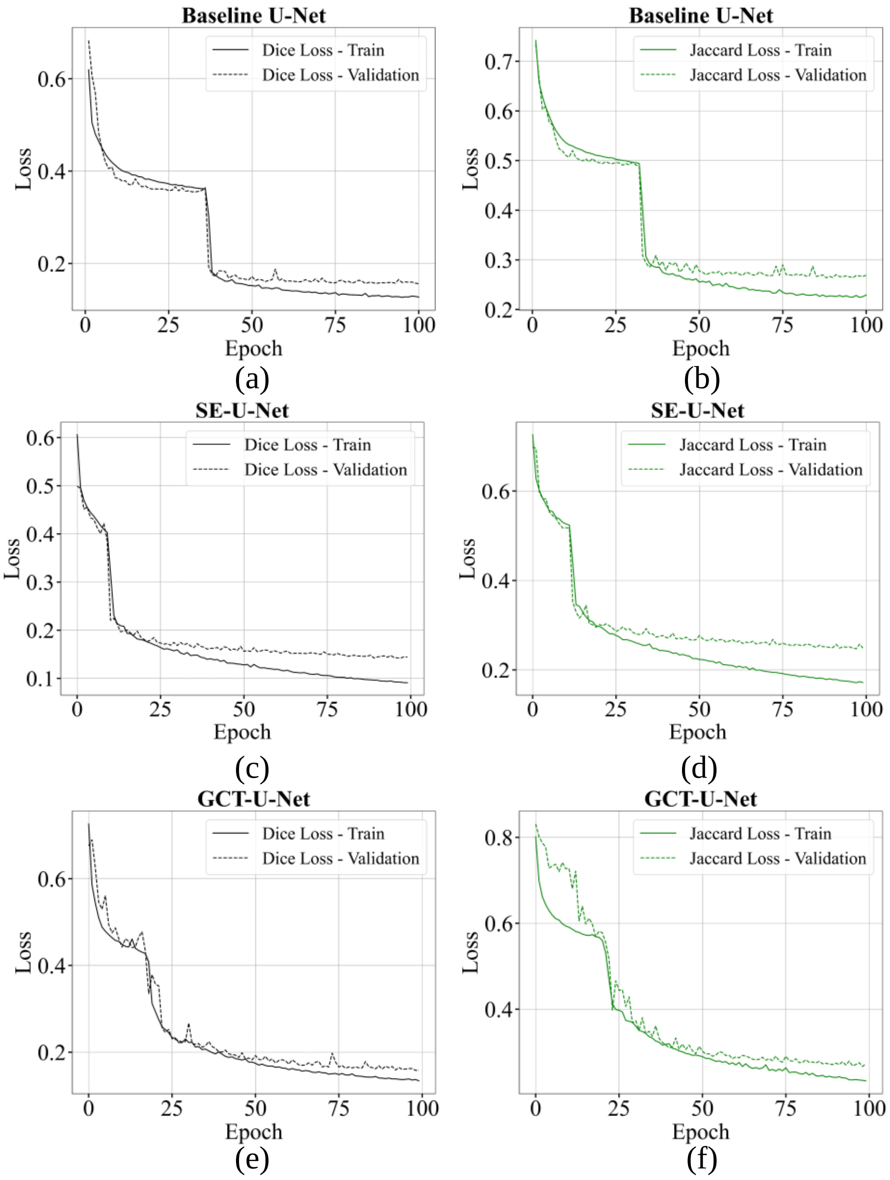


Fig. S5. Train and validation loss curve with respect to the number of training epochs on dice loss and Jaccard loss for combined image training (Σ+BF+SW) across three DL architectures, i.e. **(a, b)** Baseline U-Net, **(c, d)** SE-U-Net, **(e, f)** GCT-U-Net (csPWS-seg).

Figures S4 and S5 represent the training and validation loss curves for three different DL architectures - Baseline U-Net, SE-U-Net, and GCT-U-Net (csPWS-seg) - trained on combined Σ, BF, and SW image datasets. Each sub-figure (a-f) displays the convergence pattern while training for different epochs for the focal loss and binary cross entropy (BCE) loss (Fig. S4), as well as dice loss and Jaccard loss (Fig. S5). The GCT-U-Net, which is equipped with a Gated Channel Transformations [8] attention mechanism [9], shows the best overall performance. The training losses and the validation losses are closely aligned and decrease smoothly (shown in Fig. S4 (e)) when the model was trained with focal loss. This alignment indicates that the GCT-U-Net architecture is robust and generalizes well on unseen data, likely due to the effective channel-wise feature recalibration provided by the GCT mechanism.

#### 2.3 *Training and validation performance of csPWS-seg model on distribution-based loss functions for individually trained csPWS data*

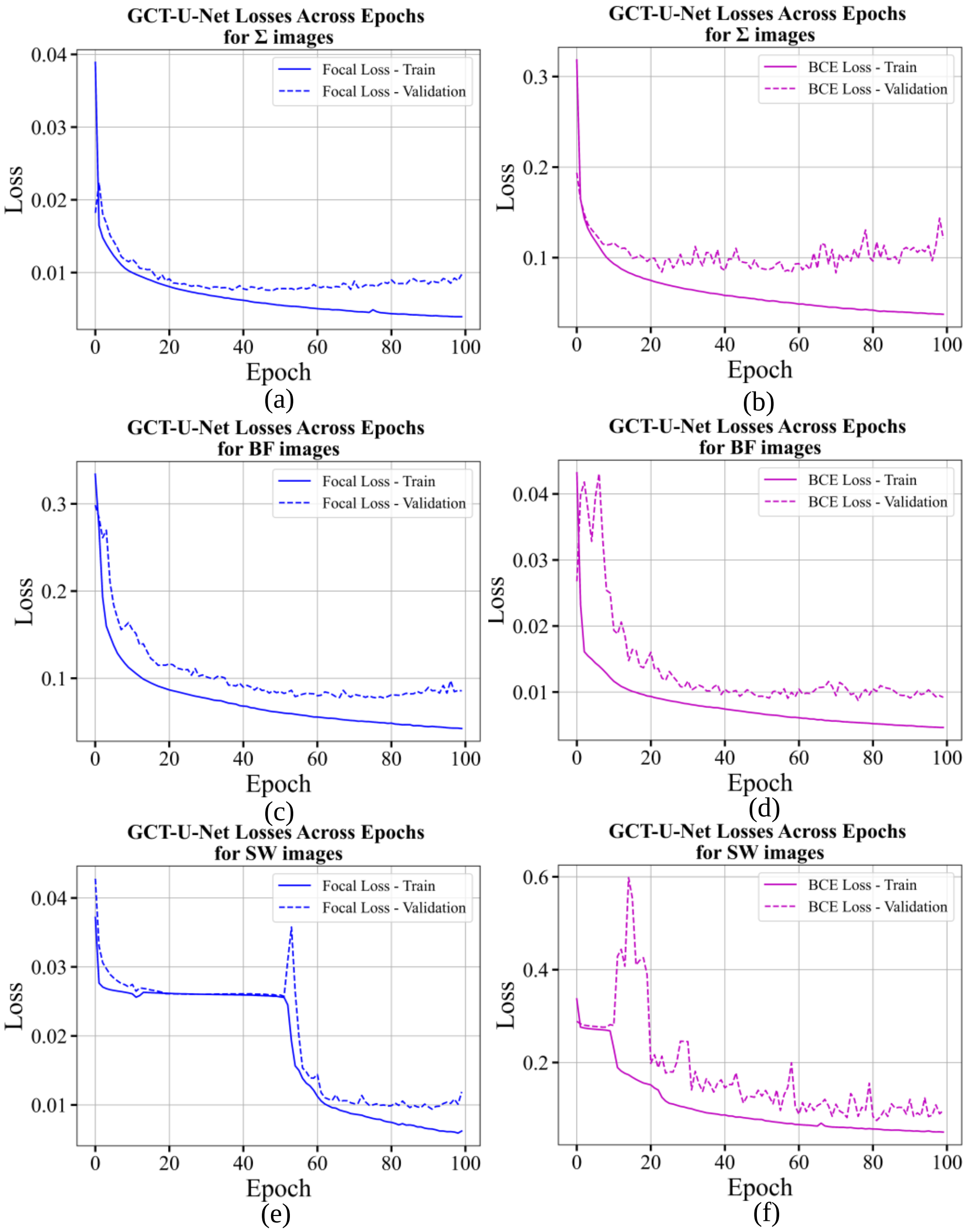


Fig. S6. Training and validation loss curves for GCT-U-Net (csPWS-seg) using focal loss and binary cross entropy (BCE) loss for separately trained feature images. **(a, b)** Σ Images, **(c, d)** BF Images, and **(e, f)** SW Images.

Figure S6 presents the training and validation loss curves for the csPWS-seg (GCT-U-Net) model across different feature image datasets, i.e., when trained separately on Σ, BF, and SW images, utilizing both focal loss and BCE loss functions. For Σ images ((a) and (b)), as well as BF images ((c) and (d)), the focal loss demonstrates a steady decrease in loss over epochs, indicating effective learning and model stability. In contrast, the BCE loss shows more variability in the validation loss, suggesting potential overfitting issues. For SW images ((e) and (f)), focal loss appears to manage fluctuations better, maintaining a more consistent validation loss curve, while the BCE loss indicates considerable variance, especially in the initial training phase. These plots underscore the varying impact of loss functions on model training and validation performance, reflecting how each loss function copes differently with the distinct characteristics of Σ, BF, and SW images.

#### *2.4 Training and validation performance of csPWS-seg model on region-based loss functions for individually trained csPWS data*

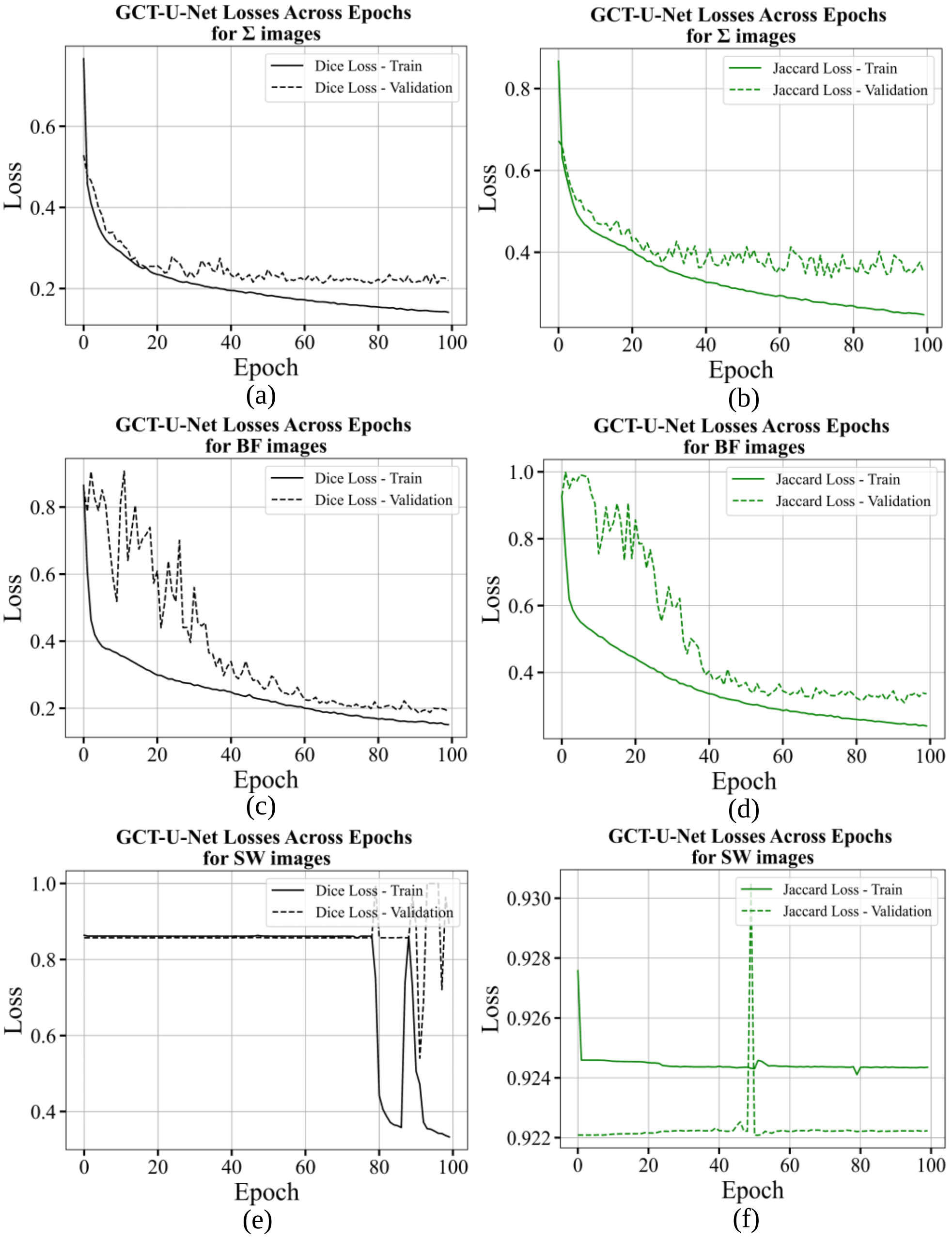


Fig. S7. Training and validation loss curves for GCT-U-Net (csPWS-seg) using dice loss and Jaccard loss for separately trained feature images. **(a, b)** Σ Images, **(c, d)** BF Images, and **(e, f)** SW Images.

Figure S7 depicts training and validation loss curves for the csPWS-seg (GCT-U-Net) model across separately trained Σ, BF, and SW images on dice loss and Jaccard loss functions. Initially, for Σ images ((a) and (b)), there is a marked reduction in both training and validation losses, demonstrating effective learning early on, which then stabilizes. For BF images ((c) and (d)), the loss curves initially show some fluctuations but gradually stabilize after about 50 epochs, indicating some challenges that are progressively overcome during training. In contrast, the SW images ((e) and (f)) display an unusual pattern, where both training and validation losses remain almost constant across all epochs, suggesting that the model does not effectively learn from this SW data.

#### *2.5 Additional performance analysis of the csPWS-seg trained with focal loss*

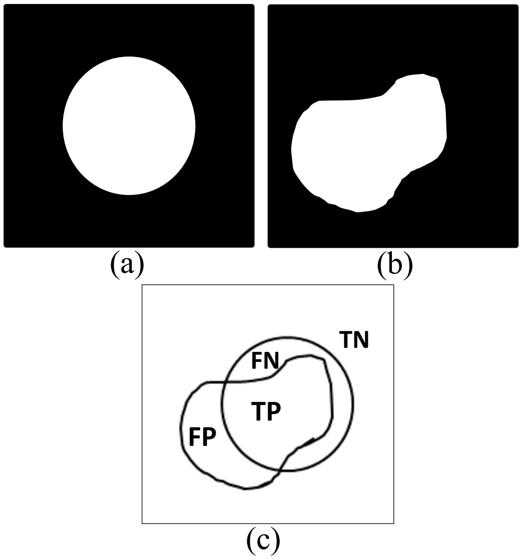


Fig. S8. Segmentation analysis. (a) Ground truth ROI (white region). (b) Predicted ROI (white region). (c) Definition of True Positive (TP), False Positive (FP), True Negative (TN), and False Negative (FN).

Table S1**.** Segmentation Analysis Metrics for csPWS-seg Trained with Focal Loss

| **Train Data** | **Test Data** | **FPR** | **FNR** | **TPR** | **TNR** |
|---|---|---|---|---|---|
| Σ+BF+SW | Σ | 0.01645 | 0.10875 | 0.89120 | 0.98355 |
| Σ | Σ | 0.01516 | 0.18355 | 0.80645 | 0.98483 |
| Σ+BF+SW | BF | 0.01753 | 0.09475 | 0.89012 | 0.98246 |
| BF | BF | 0.01846 | 0.21142 | 0.78858 | 0.98153 |
| Σ+BF+SW | SW | 0.02126 | 0.11868 | 0.89132 | 0.97873 |
| SW | SW | 0.02621 | 0.18042 | 0.81958 | 0.97381 |

Table S1 shows the performance metrics with different training and testing scenarios for the csPWS-seg trained with focal loss. The metrics False Positives (FP), False Negatives (FN), True Positives (TP), and True Negatives (TN) values are calculated for all test images using the overlap and differences between the ground truth ROI and the predicted ROI as defined in Fig. S8. FP are calculated as the sum of all instances where the predicted probability exceeds the threshold while the actual label is 0. The FPR is then determined as the ratio of FP to the sum of FP and TN. Similarly, the FN are evaluated as the sum of all cases where the predicted probability is below the threshold but the actual label is 1. The FNR is calculated as FN divided by the sum of FN and TP. The TP are the totalized cases where the predicted probability exceeds the threshold and the actual label is 1, and the TPR is determined as the ratio of TP to the sum of TP and FN. Similarly, TN are counted as the sum of all pixels where the predicted probability is below the threshold and the actual label is 0, thus, the TNR is measured as the ratio of TN to the sum of TN and FP. From the table, it is evident that the model trained on combined Σ, BF, and SW data (Σ+BF+SW), when tested with any kind of feature images results in higher TPR along with lower FNR compared to models trained on individual data types (highlighted in green) reflecting superior segmentation performance. This suggests a significant advantage in using a mixture of diversified feature images in training datasets, which enhance the model's ability by learning a wide range of characteristics features from these diverse example images. For individually trained models, Σ image only trained model shows the lowest FP when tested with the same type of images (Σ image). The SW test images, on the other hand, when tested on combined trained models, have the highest TPR values, showing the advantage of SW feature images for csPWS nuclei segmentation analysis.
